# RLKdb: A comprehensive curated receptor-like kinase family database

**DOI:** 10.1101/2023.12.18.572263

**Authors:** Zhiyuan Yin, Jinding Liu, Daolong Dou

## Abstract

Dear Editor,

Since the first plant receptor-like kinase (RLK) gene *ZmPK1* was cloned from *Zea mays* in 1990 (Walker & Zhang, 1990), this large gene family has been extensively studied and shown to play crucial roles in growth, development, and immunity (Tang *et al*., 2017). RLKs are widespread in the plant kingdom, while the biological functions of most RLKs remain largely elusive (Dievart *et al*., 2020). Given RLKs share a conserved monophyletic RLK/Pelle kinase domain, RLKs in several model plants are classified into distinct families by extracellular domains (ECDs) (Shiu & Bleecker, 2001). However, independent domain shuffling in specific lineages drives the origin of novel families, which raises a question: how about the landscape of RLKs in the whole plant kingdom? Previously, sequence homology-based methods have been widely used for RLK identification and classification, which probably will miss the distantly related proteins but with similar structures and potential novel families unmentioned in the literature. The academic community urgently requires a dedicated database for a systematic overview of the RLK gene family, providing data support for in-depth research on RLK genes. Here, we used a topology-based method to accurately isolate the RLKomes from proteomes. The obtained RLKomes were further classified into (sub)families based on ECD domains. We constructed a comprehensively curated plant RLK database (https://biotec.njau.edu.cn/rlkdb/), which contains valuable resources for investigating the origin and evolution of the RLK family and multiple online tools for personalized analysis.

## Identification of plant RLKs

To obtain the landscape of RLKs in plants, we collected 300 plant genomes with chromosome-level assemblies for identification of RLKs. In addition to some significant model species, including Arabidopsis, rice, and maize, these plant genomes encompass representatives from 4 phyla, 12 classes, and 45 orders (Figure 1A and Table S1). We adopted a previously described pipeline developed by our group to identify plant RLKs (Yin *et al*., 2023). In *Arabidopsis thaliana*, our pipeline identified 468 RLKs, representing a 72% in increase compared to the Ensembl annotation (Martin *et al*., 2023). We further examined the reliability of our pipeline with reference to the 610 putative RLKs reported by Shiu and Bleecker (2001). Among these, we observed that our pipeline missed 144 putative RLKs while predicting 2 novel RLKs. In the missed RLKs, 16 putative RLK gene model were removed from the current genome assembly, and 128 putative RLKs do not have a transmembrane domain. Collectively, our pipeline has high sensitivity and specificity and is suitable for high-throughput identification of RLKomes. In total, 220,038 RLKs were identified from 300 plant genomes. The RLKome size ranges from 1 to 2,459, with an average proteome percentage of 1.35% (Figure 1B and Table S1).

**Figure 1.**
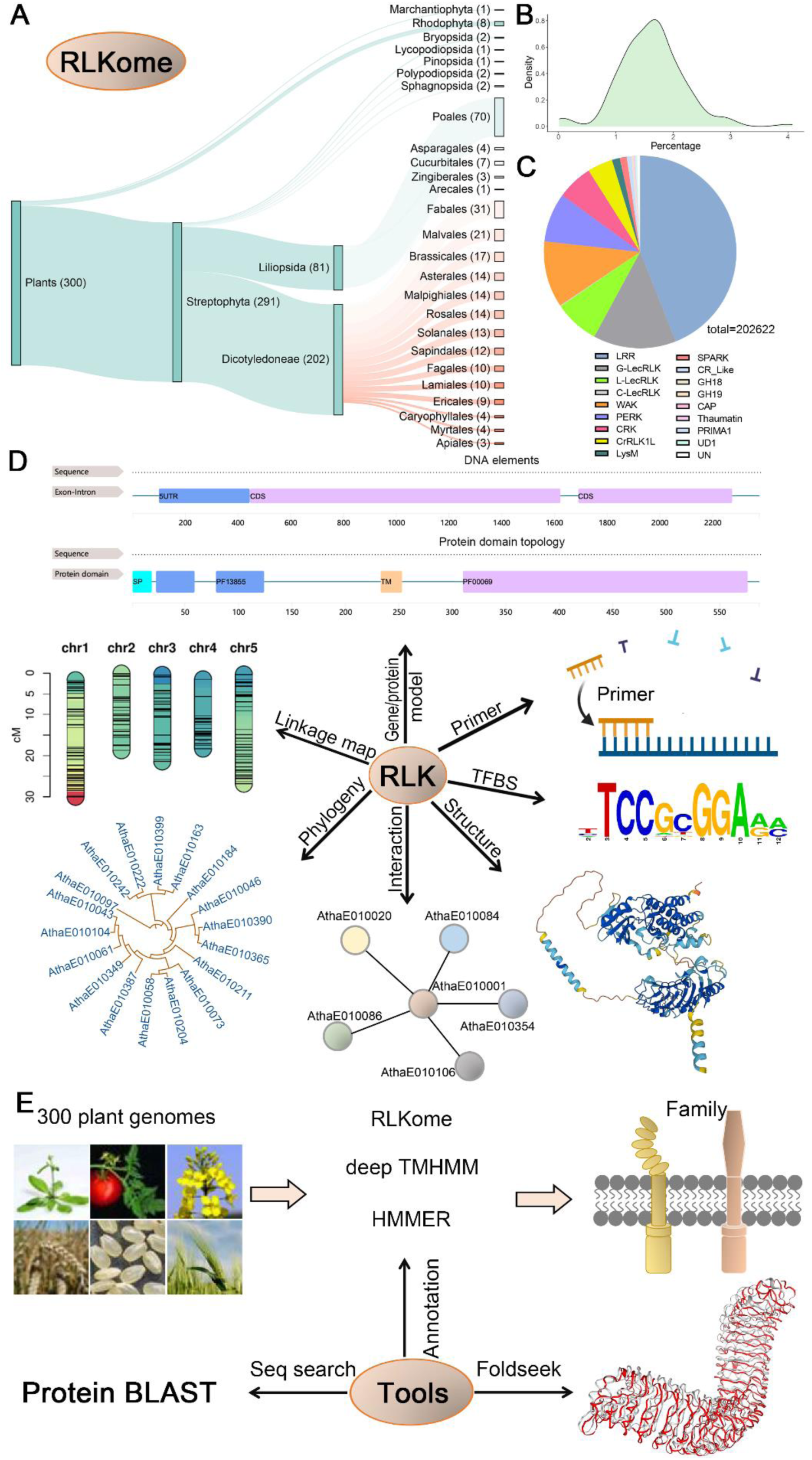
Overview of RLKdb and its functions. **(A)** Distribution of plant genomes annotated in this study. **(B)** RLKome proportions in the proteomes. **(C)** The composition of 19 RLK families in RLKdb. **(D)** Detailed information of an RLK, including chromosomal location, gene/protein model, primers, transcription factor binding site (TFBS), protein structure predicted by Alphafold2, and interaction network, etc. **(E)** Online tools for RLK/RLKome annotation and search.

## Classification of RLKs

In the past three decades, more than a dozen of RLK families have been described (Dievart et al., 2020), but a rapid and accurate pipeline for the classification of RLKome is still lacking. According to their distinct extracellular domain structures, RLKs were divided into 18 families. Among them, 15 families have known Pfam annotations. The remaining unannotated RLKs were clustered by protein sequence similarity, which further yielded the PERK and UD1 families. All the unclassified RLKs were defined as the UN family. LRR (44.0%), G-LecRLK (13.9%), and WAK (11.1%) are the largest families, which make up 69% of the RLKdb (Figure 1C). The large and well-known families occur in almost all the 300 plant genomes here, while the Thaumatin, GH19, CAP, and PRIMA1 families are only found in specific lineage.

## Web interface of the database

RLKdb has a very concise and user-friendly web interface. Through the home page or the navigation menu, users can open an RLK family (Figure S1) or RLKome page (Figure S2) to explore the database.

In the RLK family page, the first section contains the family description, its lineage coverage and a list box for switching to other families (Figure S1A). The following section is an interactive table of genomes which possess corresponding RLK family (Figure S1B). Through the load button in the table, users can load an RLK family of interests into the third section (Figure S1C). The RLK members and landscape of the family can be displayed in five panels: (1) RLK table panel shows all RLK members; (2) Linkage map panel displays the positions of RLK members in the genome; (3) Length distribution panel exhibits the distribution of RLK protein lengths; (4) Domain topology panel presents the percentage of various function domain topologies and a domain word cloud; (5) Phylogeny panel showcases the evolutionary relationships among RLK members. The RLKome page has similar layout. Its initial section provides information about the plant genome, including details on species, lineage, taxonomy, genome assembly, cultivar and more (Figure S2A). The second section is a column chart showing the number of different RLK families in the RLKome. By clicking on an RLK family name, the corresponding RLK family can be retrieved and displayed in the 5 panels that are identical to the family page.

By clicking on the hyperlinks associated with RLK IDs in the RLK table panel, users can access a dedicated RLK page displaying its detailed information (Figure 1D). In the RLK page, the first section provides a snapshot of RLK protein structure, along with essential details such as species, data source, and family information (Figure S3A). The second section contains six panels: (1) Gene model panel shows gene exon-intron structure and domain topology in protein (Figure S3B). (2) TFBS panel provides a table of transcription factor binding sites upstream (Figure S3C). (3) Primer panel offers 5 pairs of qPCR primers (Figure S3D). (4) Structure panel exhibits the 3D structure of RLK protein and its ligand binding sites (Figure S3E). (5) Interaction panel presents s a protein list that interact with RLK protein (Figure S3F). (6) Ortholog panel shows a table list of RLK orthologs in other species and the lineage coverage of the corresponding ortholog group (Figure S3G).

We also developed online tools that enable users to search and classify RLKs into different families (Figure 1E). The web-based tool allows a user to upload a proteome or transcriptome file in FASTA format (Figure S4A). The sequences undergo processing through our developed pipeline on a multi-core and GPU Linux server. For a proteome file, user will obtain an RLK annotation file containing information on signal peptide, transmembrane, kinase and other domain regions, along with an RLK sequence file. In the case of a transcriptome file, users will receive an additional ORF annotation file that highlights coding regions in the transcript sequences. To enhance database accessibility, the BLAST and Foldseek programs have been integrated to support sequence similarity and structure similarity retrieval, respectively (Figure S4B, S4C).

## Summary

In summary, we accurately annotated the RLKomes and classified RLK families of 300 plant genomes with chromosome-level assemblies. The RLKdb provides comprehensive information of RLKome, RLK family, and RLKs. We also developed an online tool for genome- and transcriptome-wide identification and classification of RLKs. The valuable resources and tools will aid evolutionary and functional studies for the community.

## SUPPLEMENTAL INFORMATION

Supplemental information is available at *Molecular Plant Online*.

## AUTHOR CONTRIBUTIONS

Z.Y. and J.L. designed and constructed the database; Z.Y., J.L., and D.D. wrote the paper.

## ACKNOWLEDGMENTS

No conflict of interest is declared.

## Supplemental Materials and methods

### Plant RLK annotation

To obtain the landscape of plant RLKome, we only focus on plant chromosome-level assemblies. A total of 300 plant genomes were gathered to establish the RLK database, comprising 153 from NCBI, 115 from Ensembl, 6 from the NGDC database, and the remaining genomes from various other public databases (Supplementary Table S1). We utilized the previously described pipeline developed by our group to identify RLKs from plant genomes. Briefly, we conducted a redundancy reduction process on plant proteomes to ensure that only the longest isoform on each gene locus was retained as the representative. Then, deepTMHMM (version 1.0.13) (Jeppe et al, 2022) was used to predict transmembrane topology, and HMMER (version 3.3.2) (Potter etal., 2018) was to identify protein kinase domains with the hidden Markov models of PF00069 and PF07714. Only the proteins were considered as RLKs if they met the criteria of having a N-terminal extracellular domain longer than 50 amino acids, one transmembrane and one C-terminal intracellular kinase domain.

### Family classification of RLKs

Despite nearly 30 years of RLK gene research, family classification has only been carried out for RLK genes in a few model species (Dievart et al., 2020). A systematic overview of the RLK gene family across the plant kingdom is yet to be conducted. Drawing from key literatures and considering the functional domain topology within RLK proteins, we have defined the following criteria for classifying RLKs into families (Supplementary Table S2).

#### (1) LRR family

The LRR-RLK family that contain various copies of leucine-rich repeat (LRR) unit in the extracellular domain (ECD) has the largest number of genes in most plant genomes (Dievart et al., 2020). The first LRR-RLK was described in 1992, when Chang et al. cloned the TMK1 (transmembrane kinase 1) gene in Arabidopsis. Given the crucial rules in plant growth, development, and immunity, LRR-RLKs have been extensively studied in the past three decades (Tang et al., 2017), such as BRI1 (BRASSINOSTEROID-INSENSITIVE1) and FLS2 (FLAGELLIN SENSING 2) that activate bacterial flagellin-induced immunity and steroid-mediated growth, respectively. The hidden Markov models (HMMs) used for the LRR-RLK classification include: PF18805, PF18831, PF18837, PF00560, PF07723, PF07725, PF12799, PF13306, PF13516, PF13855, PF14580, PF01463, PF08263, PF01462, and PF20141. Since RLKs that contain both LRR and malectin domains (malectin-like leucine-rich repeat domain and leucine-rich repeat malectin domain) are also defined as LRR-RLKs (Shiu and Bleecker, 2001), HHMs for the malectin domain (PF11721 and PF12819) are also used.

#### (2) G-LecRLK

The lectin RLKs play crucial roles in saccharide signaling and are grouped into three RLK families based on the distinct lectin domains: G-, L-, and C-type LecRLK (Vaid et al., 2013). G-LecRLK is also known as S-domain RLK, the ECD of which contains an α-mannose binding bulb-type lectin domain, a PAN (Plasminogen/Apple/Nematode), and/or an EGF (Epidermal Growth Factor) domain. The maize ZmPK1gene that was reported in 1990 is the first described plant RLK (Zhang and Walker, 1990). Other well-known G-LecRLKs includes LORE that regulates lipopolysaccharide immune sensing (Ranf, et al., 2015) and SRKs that mediate self-incompatibility (Iwano and Takayama, 2011). G-LecRLKs are also involved in plant-microbe symbiosis (Sun, et al., 2020). The hidden Markov models (HMMs) used for the G-LecRLK classification include: PF01453, PF00954, PF08276, PF00024, and PF14295.

#### (3) L-LecRLK

Plant RLKs that contain a legume lectin extracellular domain are classified into the L-LecRLK family (Vaid et al., 2013). Interestingly, functionally characterized L-LecRLKs are associated with plant immunity, while few L-LecRLKs regulate plant development or abiotic stress tolerance (Wang and Bouwmeester, 2017). Most studied L-LecRLKs are two immune receptors that sense damage-associated molecular patterns (DAMPs), including DORN1 that detects extracellular adenosine 5’-triphosphate (eATP) (Choi et al., 2014) and the receptor of extracellular nicotinamide adenine dinucleotide (eNAD+), LecR-VI.2 (Wang et al., 2019). The hidden Markov model (HMM) PF00139 was used for the L-LecRLK classification.

#### (4) C-LecRLK

C-LecRLKs are characterized by the extracellular calcium-dependent carbohydrate-binding domain (Vaid et al., 2013). Plant genomes generally possess dozens of G-LecRLKs and L-LecRLKs, while only one C-LecRLK. The C-lectin domains are mainly found in mammalian proteins that mediate innate immune responses. However, the functions of plant C-LecRLKs remain largely elusive to date. The hidden Markov model (HMM) PF00059 was used for the L-LecRLK classification.

#### (5) WAK

*Arabidopsis* WAK1 (wall-associated kinase) is the first identified WAK-RLK, the extracellular domain of which shows similarities to vertebrate epidermal growth factor (EGF)-like domains (He et al., 1996). This family plays important roles in cell expansion and pathogen resistance. Here, WAK, WAK-Like, and LRK10L (LEAF RUST10 DISEASE-RESISTANCE LOCUS RECEPTOR-LIKE PROTEIN KINASE-like) subgroups are all classified into the WAK-RLK family due to their similar extracellular domain structure (Verica and He, 2002). The hidden Markov models (HMMs) used for the WAK-RLK classification include: PF13947, PF14380, PF00008, PF08488, PF07645, PF12662, and PF12947.

#### (6) LysM

LysM-RLK is another lectin RLK family that contains extracellular LysM domains. Multiple ligands from bacteria and fungi have been shown to be perceived by the LysM domains, including chitin, β-glucans, peptidoglycans, lipopolysaccharide (Yang et al., 2022). Thus, LysM-RLKs play dual roles in plant immunity and in the establishment of the arbuscular mycorrhizal and rhizobium-legume symbioses (Zhang et al., 2021). The hidden Markov model (HMM) PF01476 was used for the LysM-RLK classification.

#### (7) CRK

Cysteine-rich receptor-like kinases (CRKs) generally contain two cysteine-rich repeat domains that are highly similar to fungal lectins (Vaattovaara et al., 2019). However, the conserved C-X8-C-X2-C motif is distinct from that in G-type lectins. This domain is also known as DUF26 or stress-antifungal domain because it has a role in salt stress response and has antifungal activity. To date, CRKs are associated with plant defense and oxidative stress responses, programmed cell death, and development (Dievart et al., 2020). The hidden Markov model (HMM) PF01657 was used for the CRK classification.

#### (8) PERK

PERKs are proline-rich extensin-like receptor kinases that constitute a small RLK family containing an extracellular proline-rich domain. Studies on the functions of PERKs are rare, most of which suggest that PERKs regulate mainly plant growth and development (Invernizzi et al., 2022). Since there are no hidden Markov model (HMM) could be used for the PERK classification, RLKs are firstly classified into different families with HMMs and the remain unclassified RLKs were clustered by protein sequence similarity. The PERKs were clustered into a sub-network.

#### (9) CrRLK1L

In 1996, Schulze-Muth et al. cloned a novel RLK in Catharanthus roseus, CrRLK1L (Schulze-Muth et al., 1996), the ECD of which was subsequently shown to be the malectin-like domain (Du et al., 2018). In recent years, CrRLKs and their peptide ligands RALF (rapid alkalinization factor) have attracted much interest. The most studied one is FERONIA (FER), which participates in a wide array of physiological processes, including cell growth and monitoring cell wall integrity, RNA and energy metabolism, and phytohormone and stress responses (Zhu et al., 2021). The hidden Markov models (HMMs) used for the CrRLK classification include: PF11721 and PF12819.

#### (10) CAP

Cysteine-rich secretory protein family (CAP) is the large family of cysteine-rich secretory proteins, antigen 5, and pathogenesis-related 1 proteins. This is an RLK family with uncommon extracellular domain. The hidden Markov model (HMM) PF00188 was used for the CAP classification.

#### (11) CR-like

The CR-like family contains an extracellular CRINKLY4 (and tumor necrosis factor receptor-like) domain, which is similar to the tumor necrosis factor receptor (TNFR) cysteine-rich region. A family member Arabidopsis ACR4 has been shown to be involved in plant development and immunity (Czyzewicz et al., 2016). The hidden Markov model (HMM) PF13540 was used for the CR-like classification.

#### (12) GH18

Glycoside hydrolase family 18 (GH18) is a type of plant chitinase. This is an RLK family with uncommon extracellular domain. The hidden Markov model (HMM) PF00704 was used for the GH18 classification.

#### (13) GH19

Glycoside hydrolase family 19 (GH19) is a type of plant chitinase. This is an RLK family with uncommon extracellular domain. The hidden Markov model (HMM) PF00182 was used for the GH19 classification.

#### (14) PRIMA1

PRIMA1 indicates the proline-rich membrane anchor 1 domain. This is an RLK family with uncommon extracellular domain. The hidden Markov model (HMM) PF16101 was used for the PRIMA1 classification.

#### (15) Thaumatin

Thaumatin-like proteins belong to the pathogenesis-related 5 (PR5) superfamily. In 1996, the RLK PR5K that contain an extracellular thaumatin-like was reported in Arabidopsis (Wang et al., 1996). This family generally has few representatives and not widespread in plant genomes. The hidden Markov model (HMM) PF00314 was used for the Thaumatin classification.

#### (16) SPARK

When Arabidopsis RLKs were firstly classified into different groups based on their extracellular domains in 2001, two were defined as unknown receptor kinases (Shiu and Bleecker, 2001). Recently, a rice RLK ARK1 (Arbuscular Receptor-like Kinase 1) was found to regulate arbuscular mycorrhizal symbiosis (Roth et al., 2018). A systematically phylogenetic analysis further showed that ARK and SIMILAR PROTEIN TO ARK 1 (SPARK1) harbor a newly identified SPARK domain in their extracellular regions, which has ancient origins in streptophyte algae (Montero et al., 2021). The hidden Markov model (HMM) PF19160 was used for the SPARK classification.

#### (17) UD1

This is a novel classified family which shares similar unknown disordered extracellular domains. The UD1-RLKs are mainly found in land plants. Plant RLKs are firstly classified into different families with HMMs and the remain unclassified RLKs were clustered by protein sequence similarity. The UD1-RLKs were clustered into a sub-network.

#### (18) UN

Plant RLKs that could not be classified by HMM search and protein sequence similarity clustering are defined as unclassified RLKs.

### Identification of RLK ortholog groups

In addition to classifying family based on functional domains, we also conducted ortholog classification based on protein sequence similarity across the 300 plant genomes. The longest isoform on each gene locus were selected to represent RLK gene. We employed Diamond-0.9.22 (Buchfink et al., 2015) for sequence alignment, Mcl-14.137 (Enright et al., 2002) for clustering, and OrthoFinder-2.3.3 (Emms and Kelly, 2019) to identify ortholog groups.

### Prediction of transcription factor binding sites upstream RLKs

To obtain transcription factor binding sites on RLK genes, the upstream 2000 bp of RLK genes were designated as candidate regions for TFBS (Transcription Factor Binding Site) search. Subsequently, curated profiles were download from the JASPAR database (Castro-Mondragon et al., 2022) to search for TFBSs using the “searchSeq” function in TFBSTools (Tan and Lenhard, 2016). Only the predicted TFBS with a minimum score of 1 were retained.

### Prediction of ligand binding sites in RLKs

To acquire the 3D structures of RLKs, we retrieved all protein sequences from the AlphaFold database (Varadi et al., 2022). Subsequently, we utilized BLASTP to identify the best matches for RLK proteins in the AlphaFold database (with an E-value threshold of <1e-10). Among the proteins that matched, over 60% shared identical protein sequences with their corresponding RLKs. We then employed the 3D structures of these proteins, identified through RLK hits, for ligand binding site prediction using p2rank (Krivak and Hoksza, 2018).

### Prediction of interaction proteins with RLKs

We employed an Interolog-based approach to forecast the proteins that interacted with RLKs. In a nutshell, we initiated the process by retrieving all experimentally confirmed interacting proteins from the STRING database (Szklarczyk et al., 2023). Subsequently, we aligned the proteome sequences of each plant species with the STRING proteins using BLASTP, employing an E-value threshold of 1e-20 and ensuring a query coverage of at least 80% and an identity of 20%. If two proteins within a proteome matched with distinct pairs of known interacting proteins from the STRING database, we inferred that these two proteins were interacting with each other. Finally, we calculate the interaction score using the following formula:

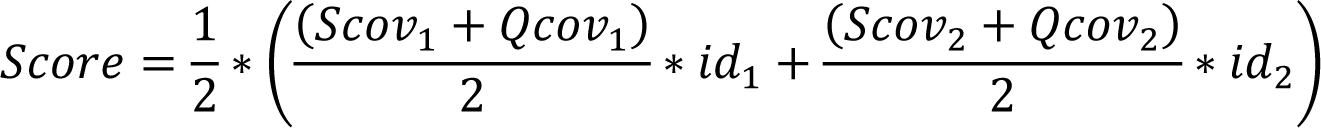

Where “Scov1” indicates the alignment length percentage in comparison to the first protein in a known interacting protein pair, whereas “Qcov1” represents the alignment length percentage relative to the querying protein. In this context, “id1” denotes the identity value between two sequence alignments. On the other hand, Scov2, Qcov2, and Id2 mirror the alignment characteristics of the second sequence in known interacting protein pairs. A higher score indicates a higher likelihood of interaction, and conversely, a lower score suggests a lower probability of interaction.

---

**Supplemental Figure 1.**
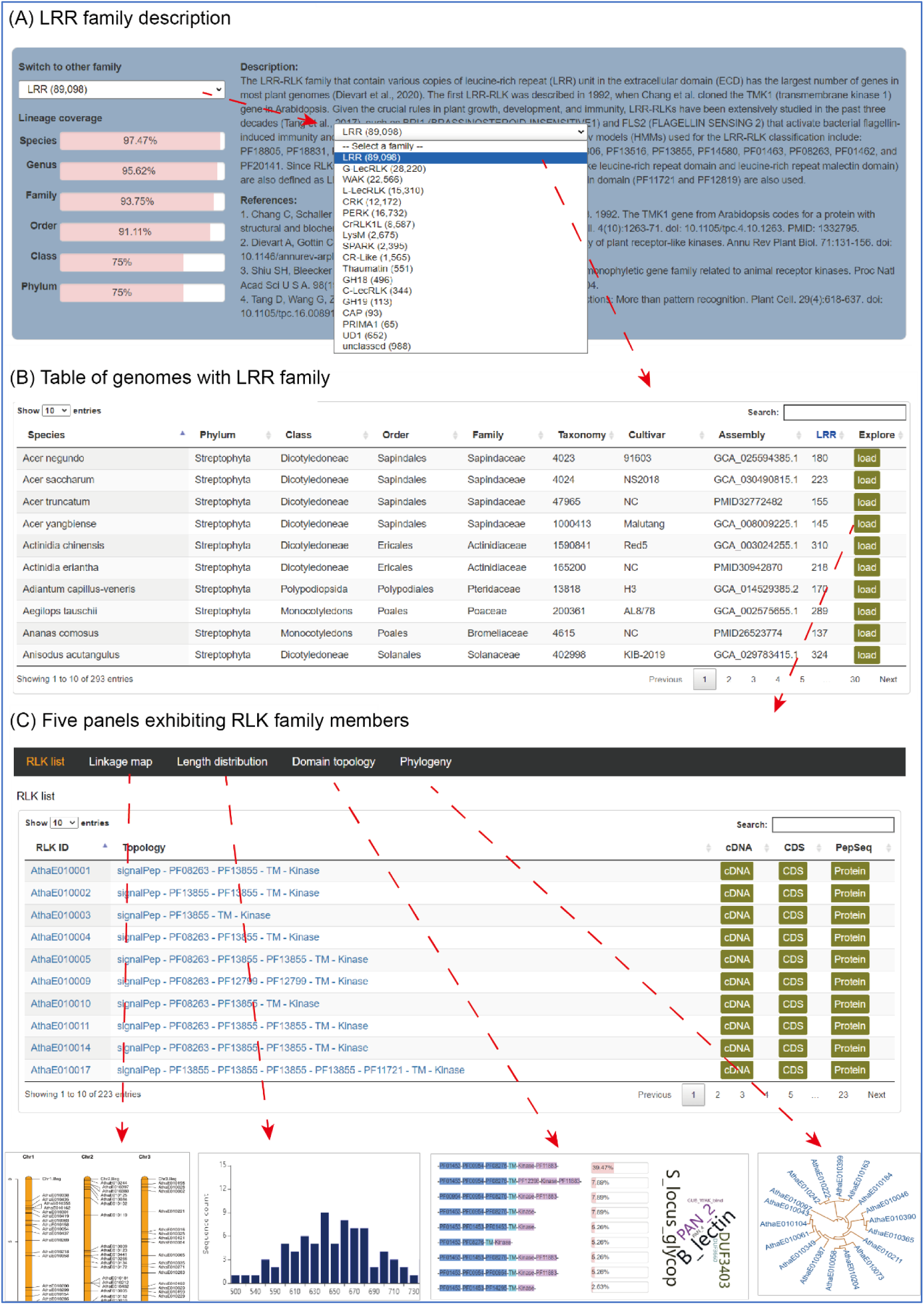
Introduction of RLK family page. **(A)** Description of RLK family. First, there is a list box for switching to other family. The family’s lineage coverage is displayed below the list box. For example, more than 97% species possess LRR family genes. The right area is the family description and related references. **(B)** Genome table. All genomes with the specified family are listed in the table with a “load” button. By clicking on the button, the RLK family in the corresponding genome can be loaded in 5 panels below. **(C)** Five panels exhibiting RLK family. All RLK members are listed in an interactive table. In addition to the functional domain topology, three buttons are provided here for downloading cDNA, CDS and protein sequence. By clicking on the hyperlinks associated with RLK IDs, users can open RLK page showing the detailed information of RLK. The second panel is linkage map. The genomic positions of RLK members are marked in the chromosomes. The third panel contains a column chart visually exhibiting RLK protein length distribution. The forth panel provides the percentages of each functional domain topology and domain word cloud. The fifth panel offers an evolutionary tree showing phylogenic relationship among family members.

**Supplemental Figure 2.**
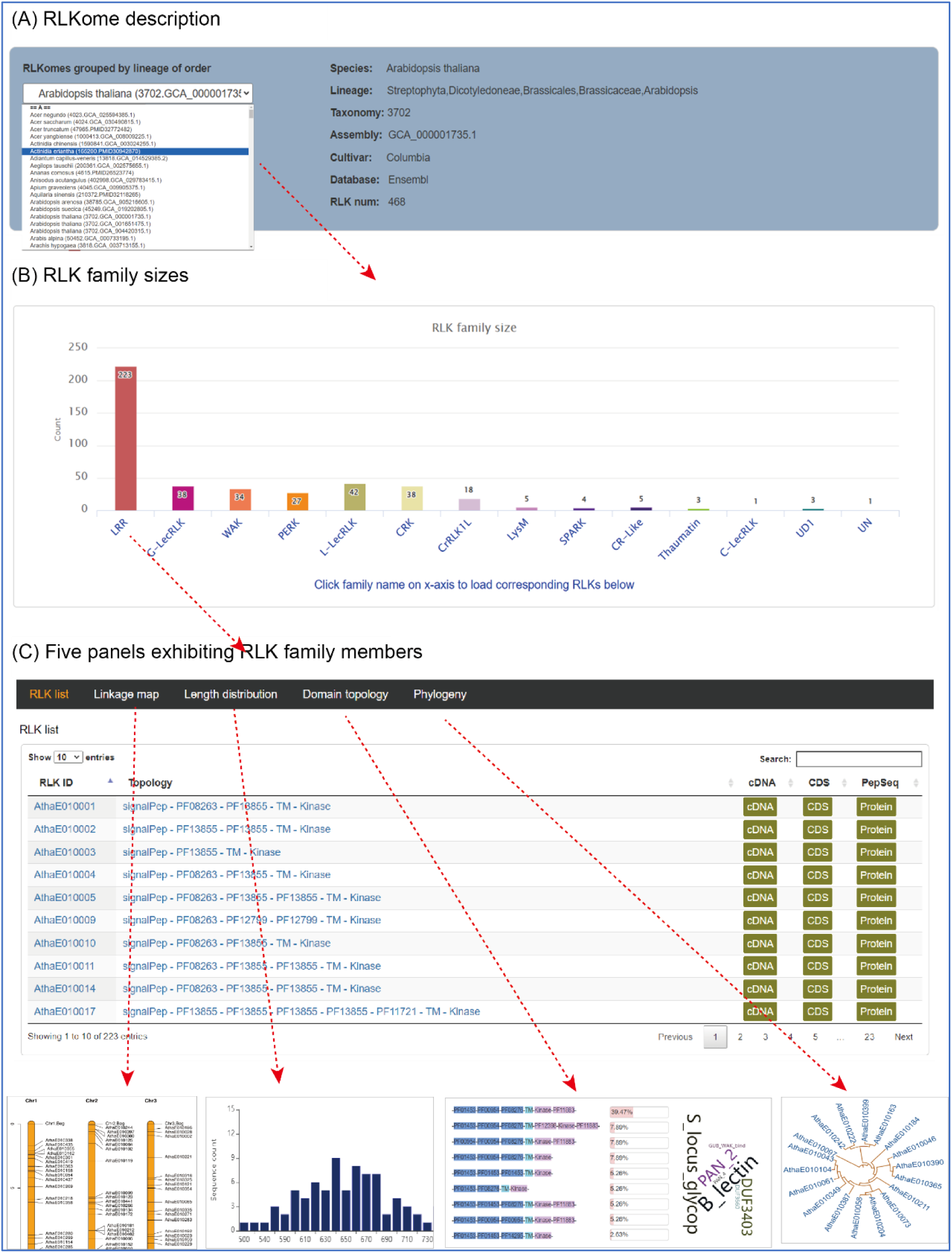
Introduction of RLKome page. **(A)** Description of RLKome page. The left area is a list box for switching to other RLKome. The right area shows RLKome information including corresponding species, lineage, taxonomy, genome assembly accession number, cultivar, database and total RLK number. (B) Column chart of family sizes. An interactive column chart exhibits RLK family size. By clicking on the family name on x-axis to load corresponding RLK family into the five panels below. (C) Five panels exhibiting RLK family. Same as S1C.

**Supplemental Figure 3.**
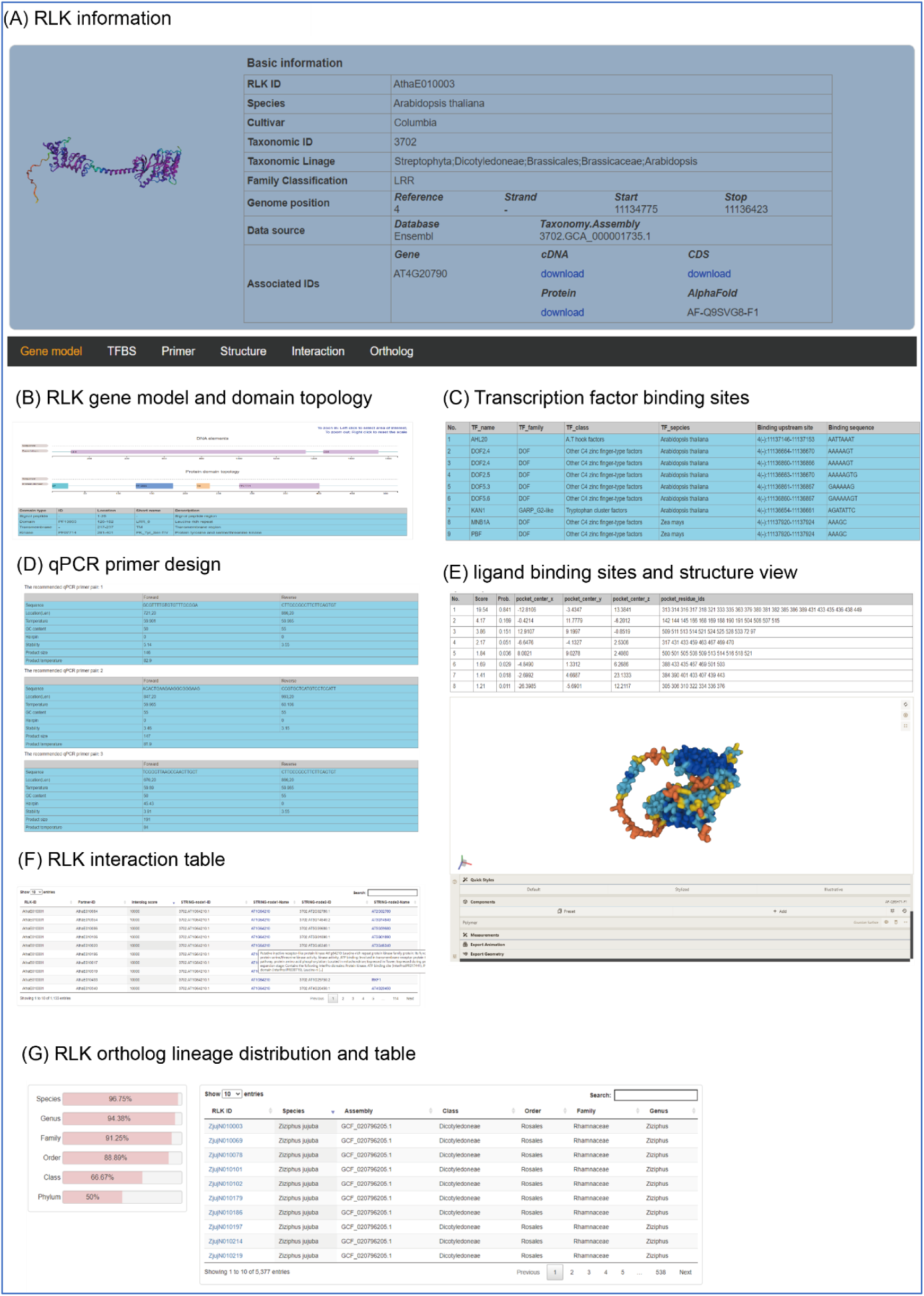
Introduction of RLK page. **(A)** RLK basic information. The left area has a snapshot of RLK 3D-strucuture. The right area is a table showing RLK basic information including RLK ID, species, cultivar, taxonomy, lineage, family, genomic position (chromosome, strand and coordinate), data source (original database and genome assembly accession number), associated sequences such as cDNA, CDS, protein and so on. The more information is divided into 6 panels. (B) Gene model panel. It exhibits gene exon-intron structure and domain topology in protein sequence. What’s more, there is a table below showing domain details including Pfam ID, location, short name and description. (C) TFBS panel. There is a table showing transcription factor binding sites upstream RLK gene. The table contains TF name, TF family, TF class, species of evidence source, TFBS and binding sequences. (D) qPCR primer panel. This panel provides 5 pairs of recommended qPCR primers. In addition to primer sequences, other information such as temperature, GC content, stability, product size, also are included in the tables. (E) Ligand binding sites and structure view panel. This panel gives a table showing possible pocket position in protein sequence. Furthermore, a 3D-structure viewer is equipped here for users to explore ligand binding sites in three-dimensional space. (F) RLK interaction panel. This panel provides an interactive table showing possible interaction proteins. A higher score indicates a higher likelihood of interaction. If mouse hover on STRING IDs, the corresponding gene function description will be presented. (G) RLK ortholog panel. The left area gives a chart showing the lineage coverage of ortholog group. In addition, the right area provides an interactive table to list all orthologs in other species. By clicking on RLK ID, users can open an RLK page showing the details of the RLK.

**Supplemental Figure 4.**
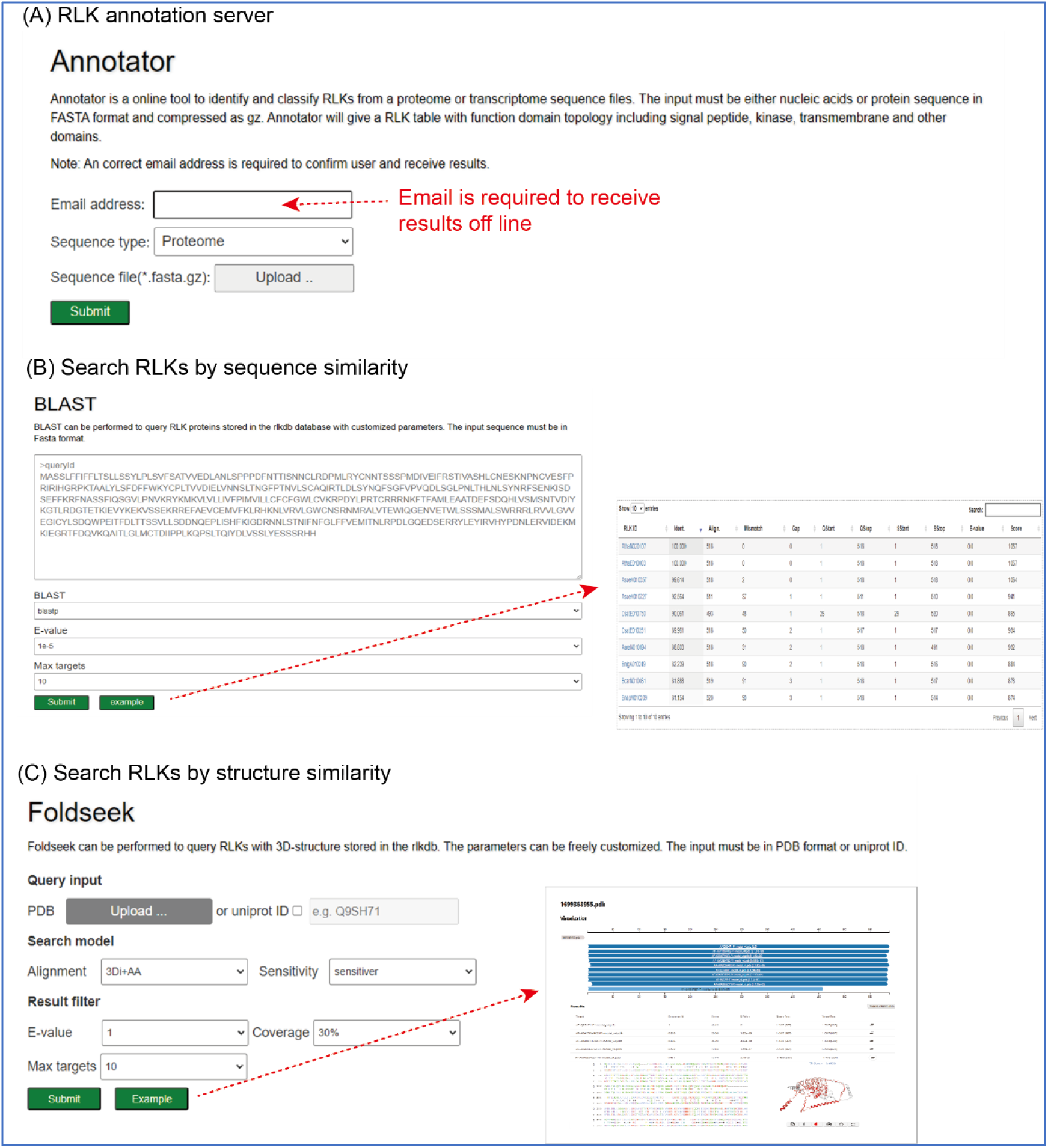
Introduction of online tools. (A) RLK annotation server. The server allows users to upload a proteome or transcriptome file in fasta format. The email address is required to receive annotation results including functional domain topology in protein and RLK family classification. For a transcriptome, the longest ORF frame in transcript sequence will be identified and translated into protein sequence. Identification of RLK will be based on the translated protein sequences. (B) BLAST tool. The tool allows users to search RLK orthologs by nucleic acids sequence or protein sequences using BLASTX or BLASTP. The queried RLKs are listed in a table below with multiple metrics such as E-value, identity, and so on. By clicking on an RLK ID, users can open the RLK page showing its details. (C) Foldseek tool. The tool allows users to search RLKs by structure similarity. Users can upload a structure file in PDB format or fill a uniport ID to search RLKs. In addition to sequence alignments, the structure alignment chart also can be exhibited.

